# A Transducing Bacteriophage Infecting *Staphylococcus epidermidis* Contributes to the Expansion of a Novel Siphovirus Genus and Implies Genus is Inappropriate for Phage Therapy

**DOI:** 10.1101/2022.10.13.512176

**Authors:** Taylor Andrews, Steen Hoyer, Karolyn Ficken, Paul Fey, Siobain Duffy, Jeffrey Boyd

**Affiliations:** Department of Biochemistry and Microbiology, School of Environmental and Biological Sciences, Rutgers University, New Brunswick, New Jersey, USA

**Author notes:** Corresponding Authors: Dr. Jeffrey Boyd.

## Abstract

The effort to discover novel phages infecting *Staphylococcus epidermidis* contributes to both the development of phage therapy and the expansion of genome-based phage phylogeny. Here, we report the genome of an *S. epidermidis*–infecting phage SEP1 and compare its genome with five other sequenced phages with high sequence identity. These phages represent a novel siphovirus genus, which was recently reported in the literature. The published member of this group was favorably evaluated as a phage therapeutic agent, but SEP1 is capable of transducing antibiotic resistance. Members of this genus may be maintained within their host as extrachromosomal plasmid prophages, through stable lysogeny or pseudolysogeny. Therefore, we conclude that SEP1 may be temperate and members of this novel genus are not suitable for phage therapy.

## INTRODUCTION

*Staphylococcus epidermidis* is a Gram-positive commensal bacterium of humans that is also an opportunistic pathogen: the most common source of nosocomial infections [1]. *S. epidermidis* infections are becoming increasingly difficult to treat due to the prevalence of antibiotic resistant strains. Instead of relying on continued development of new antibiotics [2], a promising alternative that is being approached with renewed interest is phage therapy, which uses bacteriophages to treat bacterial infections [3]. Phage therapy utilizes the mechanism of lytic phage replication to kill infection-causing bacteria. While phages can be modified or selected in laboratory conditions to optimize their performance, phage therapy relies on the diversity of naturally occurring phages of pathogenic bacteria. New phages must be isolated from the environment, characterized, and assessed for therapeutic potential [4]. However, phages that infect *S. epidermidis* remain largely under-sampled and under-studied, especially in comparison to its relative, *Staphylococcus aureus* [5].

In recent years, there has been increasing research and effort to isolate *S. epidermidis* phages [6-11]. As part of this *S. epidermidis* phage prospecting, members of a novel genus were isolated in multiple parts of the world in 2021. Pirlar et al [12] isolated, characterized and sequenced a dsDNA siphovirus, CUB-EPI_14 (ON325435.2), that is ∼43kb and has a narrow host range within *S. epidermidis*. Despite the genome of CUB-EPI_14 being labeled as likely temperate by PhageAI, which is normally disqualifying for a potential phage for therapy [13], CUB-EPI_14 lacks an integrase and the authors suggest it could be a potential candidate for phage therapy [12].

Simultaneous with the work of Pirlar et al, we also isolated a related phage from this novel genus by culturing wastewater on *S. epidermidis*, as did a third group, who has deposited their phage’s genome in GenBank without an accompanying paper (GenBank ON550478.1). We sequenced the genome of our representative (SEP1) and conducted additional host range and transduction assays. We found three additional representatives of this novel genus in GenBank (including one from an *S. epidermidis* shotgun sequencing project) and analyzed these with the three cultured phage genomes. We identified in all genomes a common phage resistance gene and a partitioning protein that is associated with being maintained in a plasmid state. Combined with our observation that SEP1 can transduce antibiotic resistance, we propose that members of this novel genus are likely temperate and therefore inappropriate for phage therapy.

## MATERIALS AND METHODS

### Wastewater Sample Screening

Aliquots of wastewater influent from a treatment plant in the mid-Atlantic United States were obtained in March 2021 and were screened for lytic phages effective against *S. epidermidis* 1457 [14]. 5mL of the wastewater samples were combined with 0.15g powdered TSB medium, 25μL of 1M CaCl_2_, and 50μL bacterial broth culture (*S. epidermidis* 1457), then incubated overnight at 37°C. A second sample of wastewater underwent the same procedure without the addition of host bacteria. After incubation, the mixtures were centrifuged at 3220xg for 15 minutes and the supernatant was passed through a 0.22 μm filter. 100μL of the filtrate was cultured with 100μL of 10^−1^ bacterial dilution of an overnight culture using the pour plate method (in 3mL of 0.3% TSB soft agar combined with 25μL of 1M CaCl_2_, vortexed and poured onto TSB 1% agar plates). The plates were then incubated overnight at 37°C and examined for the presence of plaques.

### Phage Isolation

In order to purify phages identified during the screening process, isolated plaques were picked up using a sterile glass pipette tip and the agar was deposited into a culture tube containing 2mL TSB liquid broth, 50μL bacterial broth culture, and 25μL of 1M CaCl_2_, and was incubated overnight at 37°C. This liquid culture was then centrifuged at 3220xg for 15 minutes and the supernatant was passed through a 0.22μm filter. The filtrate was diluted, and 100μL of this diluted filtered supernatant was combined with 100μL 10^−1^ bacterial dilution, 25μL of 1M CaCl_2_, and 3mL of molten TSB soft agar, vortexed, and poured onto TSB solid agar plates. The plates were then incubated overnight at 37°C and examined for plaques. This subculturing procedure was performed a total of three times to yield a purified, enriched phage stock.

### DNA Isolation and Sequencing

The DNA genome of one isolated phage was extracted using a Qiagen QIAamp MinElute Virus Spin Kit. Paired end Illumina sequencing was performed at MiGS (Microbial Genome Sequencing Center, now SeqCenter). Reads were analyzed using CPT Galaxy Phage genome assembler v2021.01 Workflow [15], which uses SPAdes Galaxy v3.12.0 [16]. This yielded three assembled contigs. These contigs were aligned manually using Aliview [17] to verify that they were identical (except for short regions of duplication due to the likely circular genomes having been assembled linearly) and to produce a complete genome without such duplications. The SEP1 genome was reoriented to mimic the linearization of relatives found using NCBI Standard Nucleotide BLAST.

### Genome annotation

The SEP1 genome was annotated using Prokka (v1.14.6, Galaxy; parameters Kingdom: Viruses, [18]. Predicted ORFs were annotated further using NCBI Standard Protein BLAST, and the sequences producing significant alignments were analyzed to determine functional gene annotations for SEP1. When the BLAST search produced multiple identical hits, we chose the annotation that was most relevant to a phage lifestyle (eg: the name given in another phage genome). Phylogenetic analysis of the SEP1 genome and other dsDNA phage genomes was performed with GRAViTy v1.1.0 (Genome Relationships Applied to Virus Taxonomy, http://gravity.cvr.gla.ac.uk/,[19].

Further functional annotation was performed. Promoter sequences were predicted by inputting the SEP1 genome into the Genome2D Prokaryote Promoter Prediction tool [20]. Rho-independent termination sites were predicted using the ARNold web tool [21]. Noncoding RNAs were found using Rfam [22]. TRNAscan-SE was used to search the SEP1 genome for transfer RNAs [23].

### Comparison to related phage genomes

To identify close relatives of SEP1, the assembled SEP1 genome was used to query NCBI Standard Nucleotide BLAST. Other phage relatives were identified by searching predicted SEP1 ORFs using NCBI Standard Protein BLAST and making note of organisms with consistent protein homology to SEP1 ORFs, whose genomes were then compared to SEP1 directly using NCBI Align Sequences Nucleotide BLAST. SEP1 and identified relatives were analyzed with GRAViTy v1.1.0 to determine their taxonomy [19]. GRAViTy results were visualized using FigTree v1.4.4 (http://tree.bio.ed.ac.uk/software/figtree/). The protein products of SEP1 predicted ORFs were analyzed to identify potential indicators of phage lifestyle, and the genomes of SEP1 and its relatives were also analyzed using the PhageAI lifestyle classifier algorithm [24].

### Host Range

The host range of SEP1 was explored via spot plating on 7 additional strains of *S. epidermidis:* 158-22, B138-22, B72-22, B76-22, B64-22, NRS101 (RP62a), and ATCC 12228. We also tested other *Staphylococcus* species isolates: *S. hominis* (160-22, B124-22), *S. haemolyticus* (B1869-21, 157-22), *S. simulans* (B149-22, B1781-21), *S. capitis* (B65-22, B1931-21), *S. lugdunensis* (B67-22, B50-22), *S. warneri* (B21-22), *S. aureus* (LAC WT, SH1000, MW2, N315). All strains are available in the collections of Jeffrey Boyd and Paul Fey.

Pour plates of each strain were prepared by combining 3mL of 0.7% molten TSB soft agar, 25μL of 1M CaCl_2_, and 100μL 10^−1^ bacterial overnight culture dilution, vortexing the mixture, and pouring it onto TSB 1% agar plates. Once the top agar solidified, 5μL of SEP1 lysate (∼10^8^ PFU/mL) was spotted onto the surface. The plates were then incubated overnight at 37°C and examined for evidence of lysis. Experiments were conducted in triplicate.

### Transduction

In order to determine whether SEP1 possesses transducing abilities, a plasmid transduction experiment was performed. A modified *S. epidermidis* strain (1457 saeR/pNF155) carrying a plasmid which is marked with an erythromycin resistance gene served as a donor strain and erythromycin sensitive *S. epidermidis* 1457 served as a recipient strain [25]. Phage-bacterial cocultures were prepared with 2mL of *S. epidermidis* 1457 saeR/pNF155 overnight culture (grown in TSB liquid broth with 10µg/mL erythromycin) was combined with 5mL TSB liquid broth, 100μL 1M CaCl_2_, and 100μL SEP1 purified phage stock. These bacteria-phage cocultures were incubated overnight at 37°C °C. The following day, SEP1 phages were harvested from the cocultures by centrifugation (13,000xg for 3 minutes) and the resulting supernatant was then filtered using sterile 0.22μm filters to remove bacterial cells. This filtered supernatant was then combined with overnight cultures of *S. epidermidis* 1457 recipient strain: 500μL *S. epidermidis* 1457 culture, 500μL TSB liquid broth, 100μL 1M CaCl_2_, and 100μL of the harvested SEP1 donor phage preparation. These cocultures were incubated at 37°C for 1 hour. Following incubation, 400μL of 1M sodium citrate was added to each, each tube was vortexed to mix, and each coculture was transferred to a microcentrifuge tube. Cells were pelleted by centrifuging at 13,000xg for 2 minutes and the supernatant was discarded. The cells were resuspended in 1mL TSB liquid broth and centrifuged again at 13,000xg for 2 minutes. The cells were then resuspended in 200μL TSB liquid broth and plated on TSB agar plates containing 10 µg/mL erythromycin and 2mM sodium citrate. As negative controls, erythromycin sensitive *S. epidermidis* 1457 was plated on erythromycin-containing TSB plates and the SEP1 phage stock was spotted onto erythromycin-containing TSB plates. Inoculated plates were incubated overnight at 37°C and were then examined for the presence of bacterial growth. The transduction experiments were performed six times.

## RESULTS

### Isolation of SEP1

Phages capable of forming plaques on *S. epidermidis* 1457 were successfully isolated from samples of wastewater influent from a treatment plant in the mid-Atlantic United States. Concentrated samples of the unenriched wastewater did produce plaques on *S. epidermidis* 1457, but the enriched wastewater produced orders of magnitude more plaques. Plating the host alone (without wastewater) did not produce any plaques. During the isolation procedure, a total of 11 plaques were harvested for potential further work. Of these isolated plaques, two were chosen for sequencing. Upon sequencing and assembly of the genomes, it was discovered that both were 100% identical and so only one (SEP1) was further characterized.

### SEP1 genome and annotation

SEP1 has a 46,473bp dsDNA genome that is likely circular (GenBank accession OP142323). It contains 72 predicted ORFs, 19 putative promoters, 1 putative noncoding RNA (which encodes a group I catalytic intron), 19 putative rho-independent terminators, and no predicted tRNAs. Several similar phage genomes were identified by BLAST (>95% identity over ≥93% of the genome), and the SEP1 genome was linearized and oriented to mimic the genomes of its close relatives, which were also isolated on *S. epidermidis* (ON550478.1, ON325435.2, Table 1).

**Table 1:** Other Proposed Members of Novel Genus

| Table 1: Other Proposed Members of Novel Genus |  |  |  |  |  |  |  |
| --- | --- | --- | --- | --- | --- | --- | --- |
| Phage Genome | Accession | Sample type | Isolation location | Length (bp) | Query cover of SEP1 | % identity with SEP1 | Phage AI Predicted Lifestyle |
| Staphylococcus phage Sazerac, complete genome | ON550478.1 | cultured isolate | IL, USA | 46428 | 94% | 96.27 | Temperate |
| Staphylococcus phage CUB-EPI_14, complete genome | ON325435.2 | cultured isolate | Germany | 46098 | 93% | 96.19 | Temperate |
| Uncultured Caudovirales phage clone 9S_3 | MF417888.1 | uncultured isolate | South Africa | 45052 | 93% | 95.53 | Temperate |
| TPA: Myoviridae sp. isolate ct5pN1 | BK030923.1 | metagenome assembled genome | USA | 46472 | 95% | 98.64 | Temperate |
| Contig of Staphylococcus epidermidis isolate Sep_B35_CVC_2019, whole genome shotgun sequence (proposed prophage) | NZ_CAJUVG01000006 | whole genome shotgun sequence from *S. epidermidis* isolate | Portugal | 46658 | 98% | 96.47 | Temperate |

SEP1’s putative protein products contain an expected assortment of phage proteins and some hypothetical proteins (Figure 1). Nine structural proteins were identified, which were similar, by BLASTp, to those of siphoviruses with long, non-contractile tails. SEP1 has both a holin and an endolysin, and 14 proteins involved in DNA replication and metabolism were identified. Two ORFs are associated with a plasmid prophage lifestyle: a parB-like protein and a potential resistance protein. The remaining 43 ORFs in the SEP1 genome are either hypothetical (34 ORFs), are identified only with a protein family or as including a known domain (9 ORFs).

**Figure 1:**
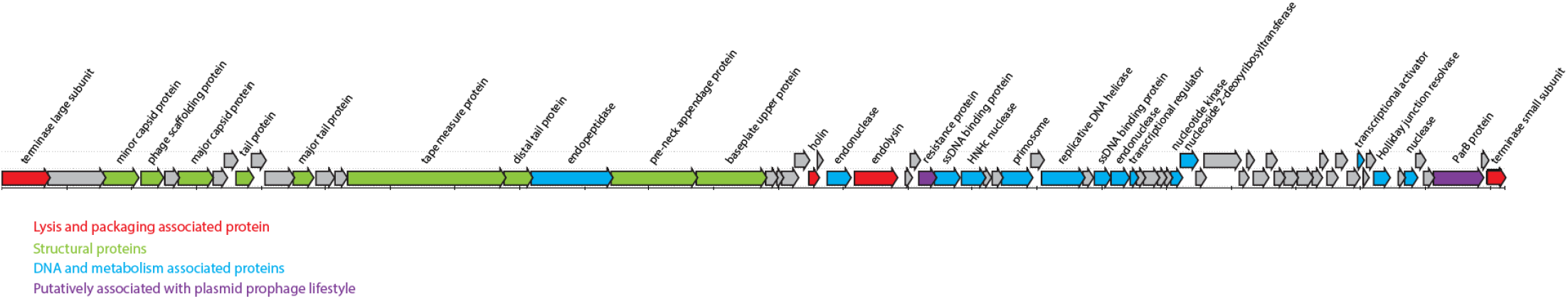
SEP1 genomic map. ORFs are annotated with predicted protein products.

### SEP1 is part of novel genus, along with other proposed members

Four other phage genomes were identified by BLASTn to be relatives of SEP1, and another by BLASTp (Table 1). Of these five relatives, only CUB-EPI_14 (ON325435.2) has yet been thoroughly described [12]. The authors of this paper note that CUB-EPI_14 appears to represent a novel genus and identifies two other potential members of genus: Uncultured Caudovirales phage clone 9S_3 (MF417888.1) and TPA: Myoviridae sp. isolate ct5pN1 (BK030923.1). These two phages were also independently identified as relatives of SEP1 during our searches. The genome of another cultured phage, Sazerac (ON550478.1), was deposited in GenBank after the manuscript about CUB-EPI_14 was submitted for publication, and we propose that Sazerac is also part of this novel genus. The final relative, Sep_B35_CVC_2019 (NZ_CAJUVG010000006), was identified due to its consistent protein sequence identity to SEP1 protein products. Although Sep_B35 is catalogued in NCBI as a contig of a *S. epidermidis* whole shotgun sequence, we argue that this contig represents a full phage genome from an infected *S. epidermidis* strain. Further, since the sample of *S. epidermidis* was sequenced as a bacterial shotgun sequencing project, not labeled as a study in phage infection, we suggest that the Sep_B35 genome represents a prophage that was being maintained within the *S. epidermidis* isolate at the time it was sequenced.

Taxonomic assignment of SEP1 and its relatives confirmed that these phages represent a novel genus within the family *Siphoviridae* (Figure 2). The six genomes form a monophyletic clade, clustered near the *Sextaecvirus* infecting other Staphylococci, among other siphoviruses. There was strong support for this group forming a novel genus (Symmetrical Theil’s uncertainty correlation 0.863).

**Figure 2:**
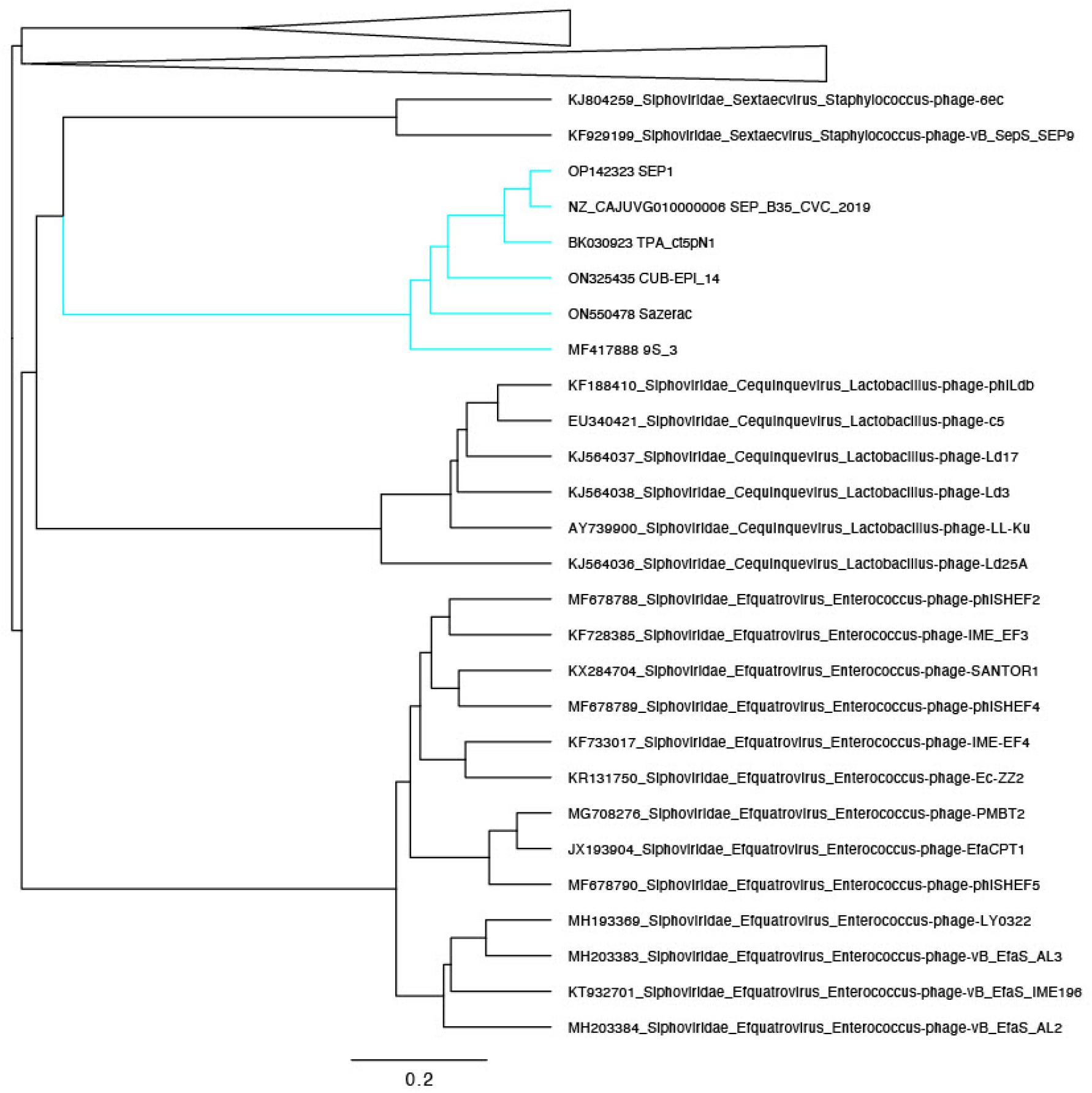
Phylogenetic tree of dsDNA prokaryotic viruses from GRAViTy, collapsed to focus on SEP1 and its relatives. Labels include GenBank accession numbers, family, order, and genus assignments, and phage names. The six genomes comprising the novel genus, including SEP1, are in the blue clade.

Following the example of Pirlar et al [12], we ran the six genomes through the PhageAI lifestyle classifier. We found that SEP1 and all members of this putative novel genus were predicted to be temperate with at least 99.95% confidence (Table 1)

### Host range results

Of the 7 *S. epidermidis* strains tested in this project, high concentrations of SEP1 was found to be capable of lysing *S. epidermidis* strains 1457, NRS101 (RP62a), B72-22, 158-22, and B138-22. Of the other *Staphylococcus* species strains tested, SEP1 was found to be capable of lysing *S. simulans* B149-22, *S. capitis* B65-22 and B1931-21, *S. lugdunensis* B67-22 and B50-22, and *S. warneri* B21-22.

### SEP1 is capable of transduction

Transduction assays were conducted three separate times and in 5/6 cases, SEP1 was capable of transducing plasmid-encoded erythromycin resistance to erythromycin-sensitive *S. epidermidis* 1457.

We were unable to identify a putative integrase gene in the genomes of SEP1 or its close relatives. ORF analysis of SEP1 and its close relatives revealed the presence of putative ParB and common phage resistance gene (Figure 1) in all six genomes (Table 2), which is partial evidence that these phages are temperate and suggests the prophages are maintained extrachromosomally.

**Table 2:** Protein Comparison

| Table 2: Protein Comparison |  |  |  |  |
| --- | --- | --- | --- | --- |
| ParB Comparision |  |  |  |  |
| Phage Genome | Original annotation | Accession | Query cover compared to SEP1 ParB | % identity with SEP1 ParB |
| Sazerac | putative adenine methyltransferase family protein | USL87160.1 | 100% | 98.20% |
| CUB-EPI_14 | DNA modification methylase | URG13538.1 | 100% | 95.99% |
| Uncultured Caudovirales phage clone 9S_3 | putative transferase | ASN69280.1 | 90% | 97.79% |
| TPA: Myoviridae sp. isolate ct5pN1 | ParB protein | DAI53229.1 | 100% | 99.80% |
| Sep_B35_CVC_2019 proposed prophage | ParB N-terminal domain-containing protein | WP_218116839.1 | 52% | 100% |
| Resistance Protein Comparision |  |  |  |  |
| Sazerac | hypothetical protein | USL87122.1 | 100% | 96.89% |
| CUB-EPI_14 | hypothetical protein | URG13503.1 | 100% | 83.23% |
| Uncultured Caudovirales phage clone 9S_3 | hypothetical protein | ASN69320.1 | 100% | 88.82% |
| TPA: Myoviridae sp. isolate ct5pN1 | resistance protein | DAI53234.1 | 100% | 100% |
| Sep_B35_CVC_2019 proposed prophage | siphovirus Gp157 family protein | WP_218116871.1 | 100% | 83.23% |

## DISCUSSION

### SEP1 is part of a novel genus

Our GRAViTy analysis strongly suggests that SEP1 and its relatives form a novel siphovirus genus. This concurs with Pirlar et al [12], who described CUB-EPI_14 as belonging to a novel genus along with 9S_3 and ct5pN1. We have also expanded this novel genus by two more phages: Sazerac and the Sep_B35_CVC_2019 putative prophage. Based on the high genetic identity between CUB-EPI_14 and the other genomes we can assume all members of this genus have long, non-contractile tails [12]. Members of this genus have already been found on three continents and we anticipate further isolates will be characterized in the upcoming years. Additional hosts may be identified for these phage as well, as we have expanded the potential hosts for members of this genus to include other *Staphylococcus* species than *S. epidermidis* (relative to Pirlar et al [12]).

### SEP1 and its relatives are not suitable for phage therapy

However, unlike Pirlar et al [12], we do not think members of this genus should be used for phage therapy. There are several characteristics that are typically screened for when assessing whether a phage could be used for phage therapy, including: host range, phage virulence, transduction potential, stability against environmental pressures, and the presence of toxin genes [26]. Bioinformatic analysis suggests members of this genus are temperate, which is contraindicated for phage therapy. Temperate phages capable of transduction have the potential to increase the pathogenicity of lysogenized bacteria by carrying virulence factors between hosts [27]. In the interest of self-preservation, prophages also typically cause lysogenized bacteria to become immune to lytic infection by other phages that share similar repression systems [26]. None of the members of this novel genus have an integrase gene, which is a key indicator of a temperate lifestyle because it allows stable integration of the phage genome into that of its host [13]. Instead, the signal that PhageAI is picking up on in the phage genomes may be the presence of t*he pa*rB gene (typically found as a *parA*-*parB* pair, implying a ParAB-*parS* system for chromosome segregation [28] and the putative phage resistance gene, which is not required in phage that only rapidly lyse their host cells [29]. Some temperate phages are known to be maintained in their bacterial host cells as extrachromosomal circular plasmids, and maintain their presence in their hosts with similar mechanisms to plasmids [29-31]. Some phage prospecting projects anticipate that these plasmid-like prophages may be isolated [32], but require that genomes have both *parA* and *parB* partitioning protein ORFs identified to be considered temperate (https://seaphages.org/forums/topic/4367/). To our knowledge, there are no characterized phage, temperate or otherwise, that only a gene for ParB, which binds to specific DNA sequences (*parS*, which vary among bacteria [33]). A BLASTp search with the ParB of SEP1 only found members of its genus and 50% coverage to other phage proteins (typically the N-terminus of ParB, data not shown). Nonetheless, we did not find a ParA homolog, which is an ATPase that assists with localization of ParB [34,35] in these six phage genomes.

The hosts of these phage, which is confirmed to be *Staphylococcus epidermidis* for four of these six phages, may offer explanation. Members of families *Streptococcaceae* and *Staphylococcacae* are known to not use a ParAB-*parS* system to ensure their own chromosome’s proper segregation into daughter cells; they use a ParB-*parS* system without a ParA [33]. Plasmids of these hosts have been found that also use a ParBS system, such as *S. aureus* plasmid SK1 [36]. Therefore, extrachromosomally maintained prophages of these hosts may also not need a ParA in order to stably vertically transmit to daughter cells. Our attempts to identify a *parS* site in SEP1 and its relatives, based on identity to *parS* sites from *S. epidermidis* [33] have been unsuccessful. However, there is low sequence identity between the ParB proteins of these phages and *S. epidermidis* and there is reason to assume that the host and phage would use quite divergent *parS* sites to bind their very divergent ParB proteins. Sequenced phages that encode both a ParA and ParB protein do not always identify a parS site [32] so we do not view our inability to find a parS site a barrier to suggesting SEP1 and its relatives temperate phage which use a ParBS system.

There is no perfect test for whether a phage is temperate and capable of creating a lysogen. We observed intermittent turbidity of plaques on *S. epidermidis* 1457 and in the spot plating experiments on other *Staphylococcus* strains, which is often considered an important phenotype of temperate phages [29]. However, turbidity can be affected by many factors [29]. Importantly, we do have affirmative experimental data for the unsuitability of members of this novel genus for phage therapy. SEP1 was capable of transducing erythromycin resistance to a previously susceptible strain of *S. epidermidis*. Transduction is a phenomenon typically associated with temperate phage, though it can be due to ‘pseudolysogeny,’ or the formation of a carrier state [37-39].

Regardless of the durability of lysogeny with SEP1, any transducing ability is empirical evidence that SEP1 and its close relatives should not be used in phage therapy.

Phage therapy remains a promising avenue of research for treating *S. epidermidis* infections, but members of this genus are not appropriate therapeutic agents. Additional isolation of *S. epidermidis* phages is needed to find obligately lytic phage.

## ACKNOWLEDGEMENTS

This work was supported by NIAID 1R01AI139100-01, NSF 1750624 and USDA MRF project NE−1028 to JMB and NSF 1453241 to SD. TPA was supported by NIGMS NIH 1T32GM139804-01. We thank the utility partner (who wished to remain anonymous) for providing the wastewater influent.

